# Co-occurring mutations in the POLE exonuclease and non-exonuclease domains define a unique subset of highly mutagenic tumors

**DOI:** 10.1101/2022.12.15.520593

**Authors:** Shreya M. Shah, Elena V. Demidova, Salena Ringenbach, Bulat Faezov, Mark Andrake, Pilar Mur, Julen Viana-Errasti, Joanne Xiu, Jeffrey Swensen, Laura Valle, Roland L. Dunbrack, Michael J. Hall, Sanjeevani Arora

## Abstract

Somatic *POLE* mutations in the exonuclease domain (ExoD) are prevalent in colorectal cancer (CRC), endometrial cancer (EC), and others and typically lead to dramatically increased tumor mutation burden (TMB). To understand whether non-ExoD mutations also play a role in mutagenesis, we assessed TMB in 447/14541 *POLE*-mutated CRCs, ECs, and ovarian cancers (OC) based on classification TMB-High (TMB-H) or TMB-Low (TMB-L). TMB-H tumors were segregated as ‘*POLE* ExoD driver’, ‘*POLE* ExoD driver plus *POLE* Variant’, and ‘*POLE* Variant TMB-H’. Intriguingly, TMB was highest in tumors bearing ‘*POLE* ExoD driver plus *POLE* Variant’ (p<0.001 in CRC and EC, p<0.05 in OC). Integrated analysis of AlphaFold2-modeled POLE models and quantitative estimate of stability indicated that multiple variants had significant impact on functionality. These data indicate that co-occurring *POLE* variants categorize a unique subset of *POLE*-driven tumors defined by ultra-high TMB, which has implications for abundance of tumor neoantigens, therapeutic response, and patient outcomes.

**Significance:** Somatic *POLE* ExoD driver mutations cause proofreading deficiency that induces high tumor mutation burden (TMB). This study defines a novel modifier role for non-ExoD mutations in *POLE* ExoD-driven tumors, associated with ultra-high TMB. These data may inform acquisition of tumor neoantigens, tumor classification, therapeutic response, and patient outcomes.

## Introduction

DNA polymerase epsilon (POLE) is an essential mediator of accurate DNA replication, based in part on its roles in DNA synthesis and DNA proofreading (1). *POLE* mutations that impair DNA proofreading lead to increased mutagenesis, and in the germline confer an increased risk of colorectal, endometrial, and other cancers (2–9). Somatic *POLE* mutations affecting proofreading are relatively rare, typically observed in ~2-8% of colorectal cancers (CRCs) and ~7-15% of endometrial cancers (ECs), and less commonly in other tumors (3). Tumors harboring *POLE* mutations that lead to proofreading defects are typically ultra-hypermutated (>100 mut/Mb) and have a specific context of mutational signatures (COSMIC signatures 10a and 10b) (10). The increased tumor mutation burden (TMB) in such tumors is typically associated with benefit from immune checkpoint inhibitor (ICI) therapy (11). Better understanding of the mechanisms by which *POLE* mutations affect TMB has significant clinical relevance for patient outcomes and treatment decisions (11–13).

Somatic mutations associated with enhanced TMB, and proofreading deficiency are typically observed as hotspot mutations in the exonuclease domain (ExoD) of POLE, such as P286R, V411L, S297F, A456P, and S459F (10), considered driver mutations. Current evidence suggests that most mutations located outside the ExoD or those leading to a truncated protein, do not have an impact on the proofreading function of the polymerase or on the tumor mutational landscape. However, *POLE* variants of uncertain significance (VUS), typically in the non-ExoD or non-hotspot regions of ExoD, are sometimes concurrent with a *POLE* ExoD driver mutation and/or microsatellite instability (MSI) (9,14,15). We hypothesized that the presence of these non-pathogenic *POLE* variants might further increase *POLE* ExoD driver-associated mutation rates. Here, we performed a retrospective analysis of 447 genomic profiles of tumors with *POLE* mutations to investigate the effect of co-occurring *POLE* non-pathogenic variants on TMB, protein stability, and clinical and molecular features.

## Results

### *POLE* variants in colorectal, endometrial, and ovarian tumors

The clinical and demographic characteristics for patients with CRC, EC, and OC genomically profiled for *POLE*, TMB, and (where relevant) microsatellite stable/microsatellite instable (MSI/MSS) status by Caris Life Sciences (CLS) are in **Supplementary Table 1.** *POLE* mutations were observed in 4.9% of CRCs (92/1870), 6.9% of ECs (307/4481), and 0.6% of ovarian cancers (OCs) (48/8190) (**Figure 1A**).

**Figure 1.**
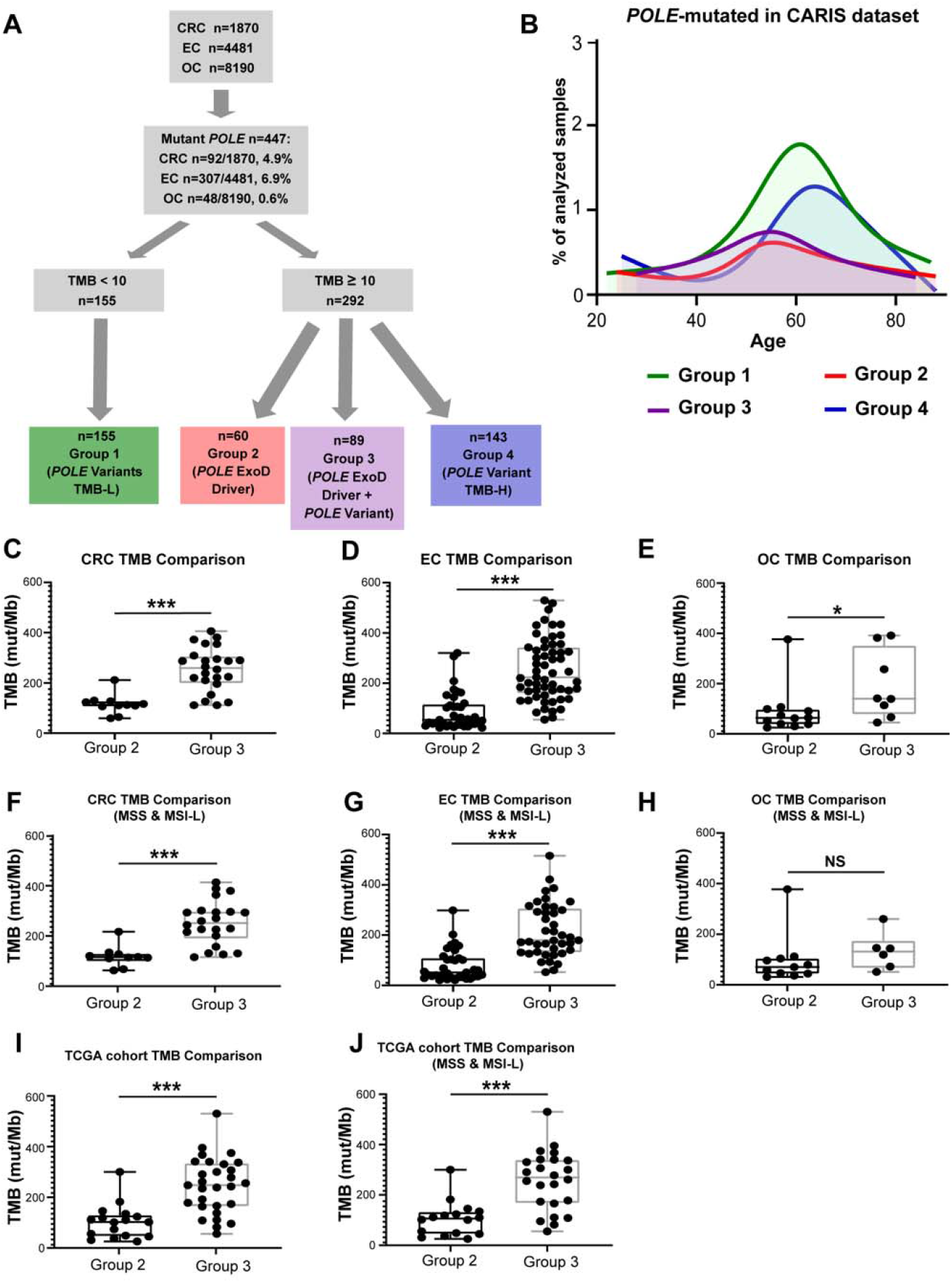
Characterization of *POLE* mutations in the CLS and TCGA dataset. **A.** Flowchart and analysis tree for CRC, EC, and OC tumors by *POLE* mutations, TMB, and microsatellite status. Among 1870 CRC, 4481 ECs and 8910 OC tumor genomic profiles, a total of 447 carried *POLE* mutations. Clinically relevant TMB cut-points were used to define the TMB-H (≥10 mutations/MB) and TMB-L (<10 mutations/MB) cohorts. *POLE* mutation cohorts along with TMB and MSI/MSS status were defined (also see Supplementary Table 1 and Supplementary Table 2). TMB-L tumors with *POLE* variants but no established *POLE* ExoD driver are referred to as ‘*POLE* variants TMB-L’ (Group 1, MSS or MSI). TMB-H tumors with *POLE* ExoD driver (Group 2), *POLE* ExoD driver plus *POLE* Variant (Group 3) and ‘*POLE* Variant TMB-H’ group (Group 4) with MSS or MSI. **B. Age distribution of patients in the CLS cohort** with *POLE*-mutated tumors (n=447) designated as Group 1 (green), Group 2 (red), Group 3 (purple) and Group 4 (blue). **C-E. mTMB comparisons between Group 2 and 3 CRCs** (**C**), **ECs** (**D**), **and OCs** (**E**). See Supplementary Table 1 and Supplementary Table 4 for more details and other comparisons. **F-H. mTMB comparisons between Group 2 and 3 genomic profiles of CRC** (**F**), **EC** (**G**), **and OC** (**H**). MSI-H tumor profiles were removed from this analysis. See Supplementary Table 1 and Supplementary Table 4 for more details. **I-J. TCGA cohort mTMB comparisons between Group 2 and 3 tumors**, in (**I**) MSI-H or MSS tumor profiles were included and in (**J**) only MSS tumor profiles were included. Due to smaller sample size per tumor type, analyses were pooled. See Supplementary Table 5 for more details. A Mann-Whitney test was performed and ***, p<0.001; *, p<0.05; NS, non-significant.

Within this dataset of tumors harboring *POLE* mutations (n=447/14541), low TMB (TMB-L, <10 mut/Mb) was observed in 39.1% of CRCs (36/92), 30.9% of ECs (95/307), and 50.0% of OCs (24/48) (**Figure 1A**, **Supplementary table 1**). TMB-L tumors had *POLE* variants but no established *POLE* ExoD driver mutations (as defined in (10) and **Methods**). This group is subsequently referred to as ‘*POLE* variants TMB-L’ (Group 1, **Figure 1A, Supplementary Table 1, and 2).** In contrast, high TMB (TMB-H, ≥10 mut/Mb) was observed in 60.9% of CRCs (56/92), 69.1 % of ECs (212/307), and 50.0% of OCs (24/48) (**Figure 1A, Supplementary Table 1**). TMB-H tumors could be segregated into three groups, subsequently referred to as ‘*POLE* ExoD driver’ (Group 2), ‘*POLE* ExoD driver plus *POLE* Variant’ (Group 3), and ‘*POLE* Variant TMB-H’ group (Group 4, lacking an established ExoD driver) (**Supplementary Table 1**, **Figure 1A**, **Supplementary Table 2)**. The Group 2 and 3 tumors typically had a single established *POLE* ExoD driver; however, five tumors had more than one (**Supplementary Table 2**). Typically, Groups 2 and 3 occurred in younger patients, while Groups 1 and 4 tumors occurred more frequently in older patients (median ages at diagnosis: 55.5, 55, 62 and 65, respectively; **Figure 1B, Supplementary Table 3**).

Interestingly, Group 3 had the highest median TMB (mTMB) compared to Groups 1 and 2, across the three cancer types (p<0.001 for CRC and EC, p<0.05 for OC, **Figures 1C, 1D, 1E, Supplementary Table 4),** even when MSI-high tumors were excluded from the analysis (significant differences (p<0.001) for CRC and EC, but not for OC, likely due to small sample size; **Figure 1F, 1G, 1H, Supplementary Table 4)**. We validated our findings by analyzing the sequencing data from The Cancer Genome Atlas (TCGA), which included 46 tumors (78% EC or CRC) with *POLE* variants (access date: Feb 2022). In this dataset, a significantly higher mTMB was observed in Group 3 (*POLE* ExoD driver plus *POLE* Variant) versus Group 2 (*POLE* ExoD driver) (p<0.001; Figure **1I**, **Supplementary Table 5**), even when excluding MSI-high tumors from the analysis (p<0.001; **Figure 1J**, **Supplementary Table 5**).

Based on the similar results obtained for the three different cancer types in the CLS dataset, we combined them to obtain better statistical power. We analyzed how mTMB changed with increasing number of *POLE* variants in the presence of a *POLE* ExoD driver (**Figure 2A** and **2B**). The mTMB significantly, and progressively, increased as the number of *POLE* variants increased; however, it stabilized and did not increase further beyond acquisition of two *POLE* variants (**Figure 2A**, p<0.001). As expected (16), P286R and V411L were the two most common *POLE* ExoD drivers across the three cancer types in the CLS dataset (67% of the tumors with a *POLE* ExoD driver had either P286R or V411L). The analysis of tumors according to the specific POLE ExoD driver (P286R, V411L or any other ExoD driver), confirmed the independent nature of the association, i.e., the increasing mTMB with increasing number of *POLE* variants, independently of the POLE ExoD driver (**Figure 2B)**.

**Figure 2.**
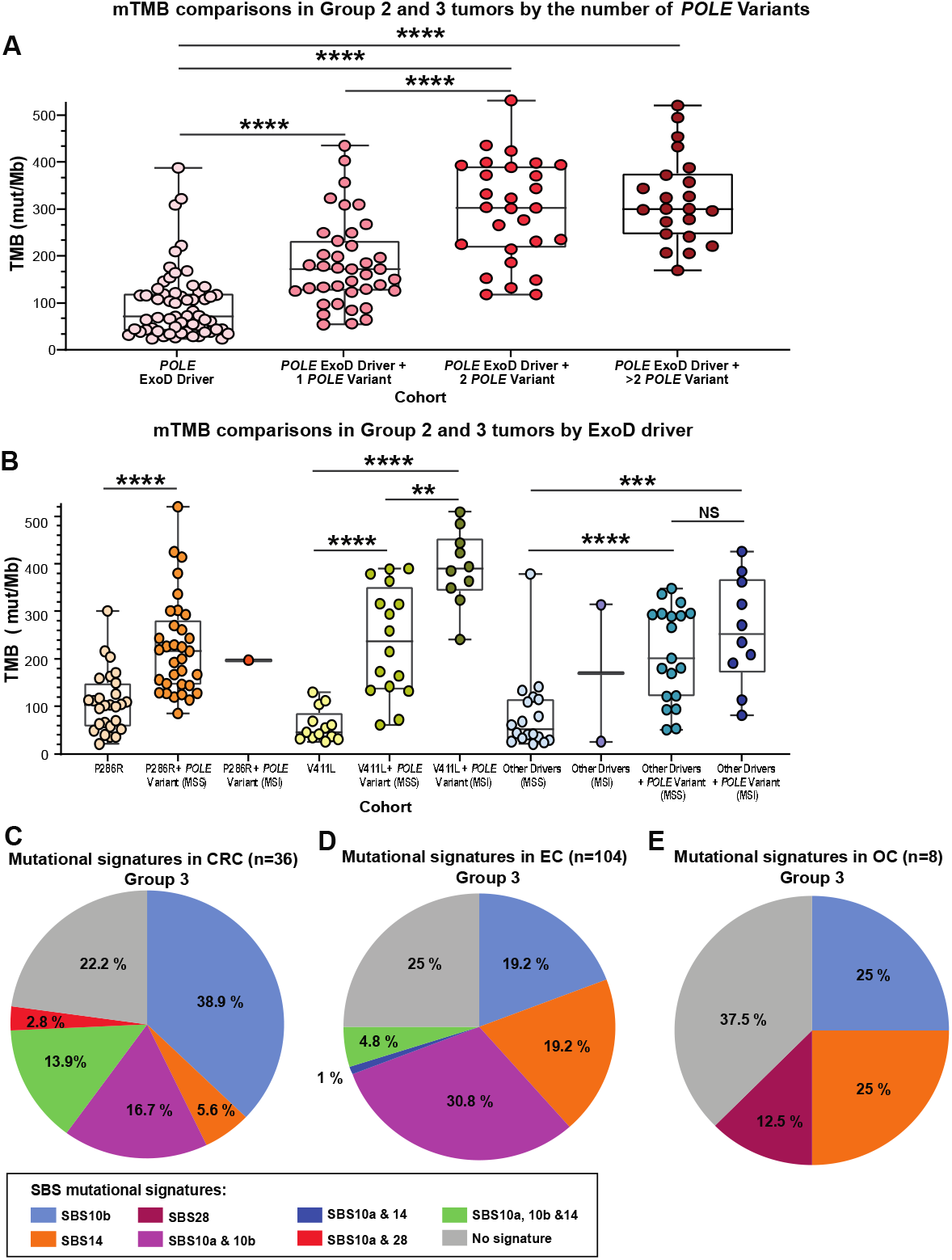
**A. mTMB comparisons in Group 2 and 3 tumors by the increasing number of *POLE* Variants.** Data are combined analysis of CRC, EC, and OC genomic profiles in the Caris dataset. Each filled round circle represents a tumor genomic profile; data are shown for Group 2 ExoD drivers, and for Group 3 by the ExoD driver plus the number of variants (1, 2, and >2 variant(s)). **B. mTMB comparisons in Group 2 and 3 tumors by ExoD driver alone (P286R, V411L, or other driver(s) combined) or in conjunction with *POLE* variants**. Each filled round circle represents a tumor genomic profile; data are shown for the driver alone and/or plus *POLE* variant. Data in ‘other drivers’ was combined due to lower numbers. The data are segregated for MSS or MSI status where relevant. A few statistical comparisons were not performed due to ≤ 2 data points. **A and B**. A Mann-Whitney test was performed and ****, p<0.0001; ***, p<0.001; **, p<0.01; NS, non-significant. Corrections for multiple comparisons were performed using the Benjamini Hochberg False Discovery Rate test. **C-E. Three nucleotide sequence context of *POLE* variants in Group 3 tumors**. All COSMIC mutational signatures associated with POLE ExoD driver defects (SBS 10a, SBS 10b, SBS 14, and SBS 28) were assessed (16). Pie chart distribution of SBS 10a, SBS 10b, SBS 14, and SBS 28 in CRC **(C),** EC **(D),** and OC (**E**). Each of these signatures has a primary mutation which has been described as a “hotspot” (16); SBS 10a is C>A in TCT context; SBS 10b is C>T in the TCG context; SBS 14 is C>A in the NCT context (N is any base); and SBS 28 is T>G in the TTT context. In addition to these primary “hotspots”, all mutations that comprise >1% of the genome signature of interest were counted, capturing 88-90% of each signature in the analysis.

### Location and nature of the *POLE* variants identified in Group 3 and 4 tumors

The *POLE* variants that accompanied the ExoD drivers (Group 3) included: 12 ExoD variants (all missense), and 143 non-ExoD variants (12 disruptive (2 frameshift and 10 nonsense), and 131 missense). It would be extremely important to analyze the 3-nucleotide context of the Group 3 *POLE* variants to elucidate if these variants might have risen as consequence of the proofreading defect. We assessed the 3-nucleotide context, according to the cancer type and overall, considering the 3-nucleotide contexts associated with COSMIC mutational signatures SBS (single base substitution) 10a, 10b, 14 and 28 for POLE proofreading defects (16,17) (**Figure 2C, 2D, 2E and Supplementary Figure 1A** (**overall**). In CRC, the primary mutation associated with SBS 10b (TCG>TTG) comprised 36.1% of *POLE* variants. TCT>TAT is considered primary for both SBS 10a and SBS 14 and comprised 13.9% of *POLE* variants (**Figure 2C**). Overall, in CRC, 77.8% of the *POLE* variants occurred in *POLE* ExoD signature sequence contexts (**Figure 2C**). In EC, TCG>TTG substitutions comprised 17.3% of *POLE* variants, and TCT>TAT comprised 4.8% of variant *POLE* variants. Interestingly, GCG>GTG, which is a minor mutation associated with both the SBS 10a and SBS 10b signatures, comprised 26.9% of *POLE* variants. This specific mutation has been previously associated with *POLE* driver mutations with defective MMR (16). CCG>CTG, which is a minor mutation in SBS 14, was also a common mutation in Group 3 tumors (10.6% of *POLE* variants). Overall, in EC, 75% of the *POLE* variants occurred in *POLE* ExoD signature sequence contexts (**Figure 2D**). In OC, there were 2 TCG>TTG mutations of SBS 10b and 2 minor mutations of SBS 14. The OC group has a small sample size: of these, 62.5% of the *POLE* variants occurred in *POLE* ExoD signature sequence contexts (**Figure 2E**). Overall, the majority *POLE* variants in Group 3 tumors occurred in the 3-nucleotide context associated with POLE defects.

The *POLE* variants in Group 4 tumors were: 27 ExoD variants (2 deletion-insertion, 1 splice site variant, and 24 missense variants) and 127 non-ExoD variants (8 nonsense mutations, 10 frameshift mutations, 5 deletions, 1 duplication, 1 insertion, 2 canonical splice site variants, and 100 missense variants). Among the MSS TMB-H subset of Group 4 tumors, *POLE* variants associated with TMB-H were mostly missense (8/14), however nonsense and other alterations were also observed (6/14) (**Supplementary Figure 1B**, see Supplementary Results).

### Effect on protein stability of the *POLE* missense variants in Groups 2, 3 and 4 tumors

To assess the effect of the POLE missense variants, human POLE structure models were generated using AlphaFold2 (18,19) (**Supplementary Methods**). The wildtype AlphaFold2 structure models generated in this study are available at https://zenodo.org/record/7395412#.Y44AwOzMJqs (DOI 10.5281/zenodo.7395412). We utilized the Rosetta ddG_monomer application (20) to predict changes in stability (or Gibbs free energy, ΔΔG) of the variants compared to the corresponding wildtype (WT) residues (**Supplementary Table 6**). The WT values are highly reproducible: for instance, the standard deviation of the mean energies for 5 runs without DNA was only 0.16 kcal/mol. Using a cutoff of 1.45 kcal/mol for significant ΔΔG, Group 2 and 3 tumors were annotated by mTMB and the number of *POLE* variants (**Figure 3A**). Group 2 tumors contained mostly destabilizing ExoD driver mutations according to Rosetta ddG_monomer (**Figure 3A**, ΔΔG ≥ +1.45 kcal/mol). In contrast, the *POLE* variants in Group 3 tumors are mostly stabilizing (**Figure 3A**, ΔΔG ≤ −1.45 kcal/mol). Further, Group 3 tumors with P286R driver generally had one or two additional *POLE* variants, where most are structure stabilizing (**Figure 3B**). Group 3 tumors with V411L driver that had the highest mTMBs tended to have multiple *POLE* variants (2 to 8) per tumor, with a range of structure stabilizing or destabilizing variants. For Group 3 tumors with other drivers, most *POLE* variants were structure stabilizing.

**Figure 3.**
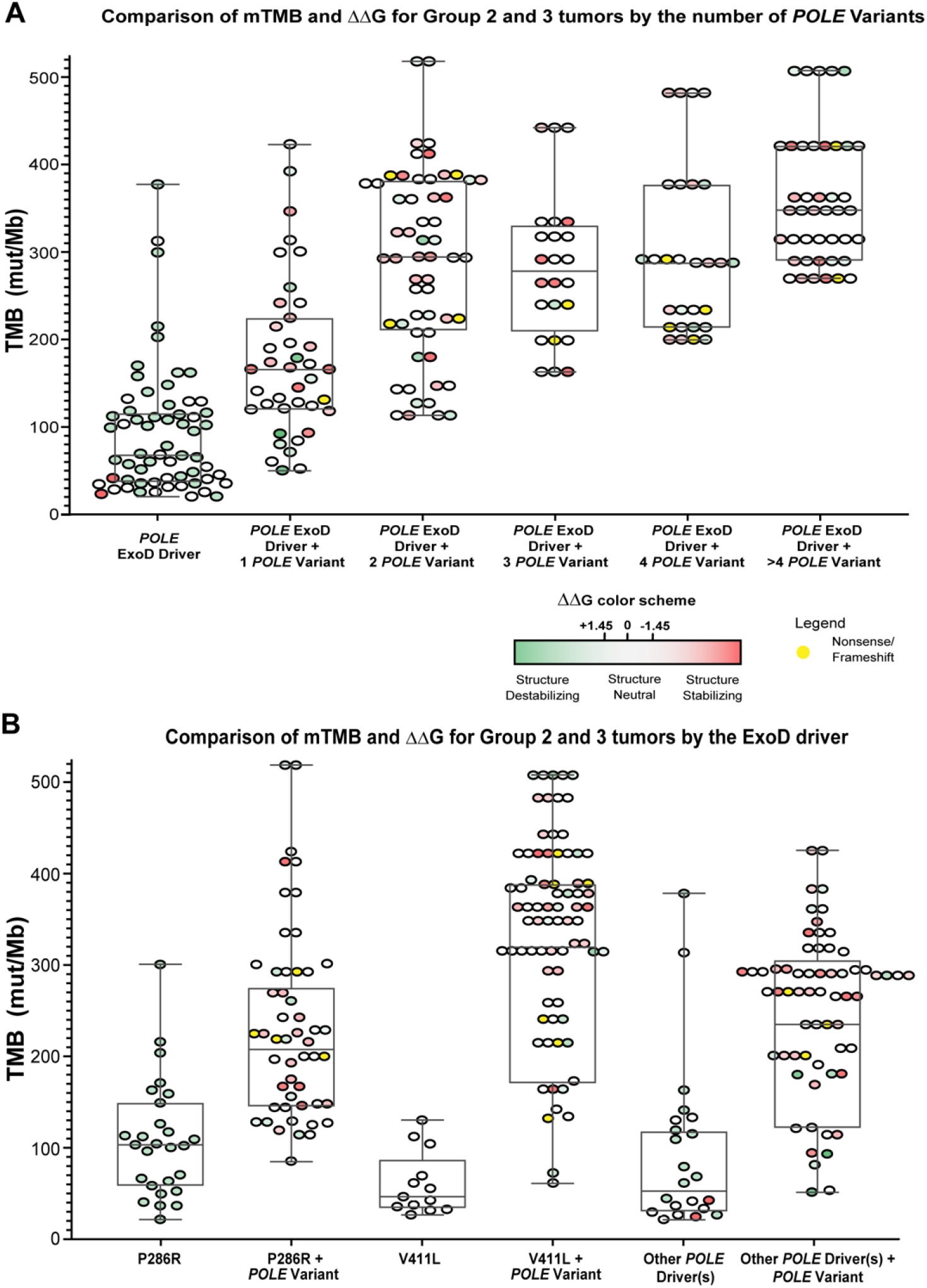
Comparison of mTMB and ΔΔG values in Group 2 and 3 tumors in the CLS dataset. With AlphaFold2 DNA unbound model and Rosetta ddG_monomer, we generated 25 repacked decoys for each mutation and compared the average energy score for these decoys to an average for 25 decoys of the wildtype protein. For mutations in the NTL, we performed calculations on both models (with and without DNA; (also see **Supplementary Figure 1**, with DNA). For those in the CTL, we performed calculations only on the DNA-unbound model. We used a cutoff of ±1.45 kcal/mol for significant ΔΔG, corresponding to ~3 standard deviations of the differences of the mean Rosetta scores for WT and mutant structures (see Methods). **A. Comparison of mTMB and ΔΔG values in Group 2 and Group 3 tumors by the number of *POLE* variants**. Data for CRC, EC, and OC genomic profiles were combined, and ΔΔG values were plotted against the mTMB. For Group 2 or 3 data with + 1 *POLE* variant plots, each filled round circle represents a single tumor genomic profile. **B. Comparison of mTMB and ΔΔG values in Group 2 and Group 3 tumors by *POLE* ExoD driver.** Data for CRC, EC, and OC genomic profiles were combined, and ΔΔG values were plotted against the mTMB. **A and B**. Group 3 tumors with multiple variants, a circle next to another circle (without any space) represents a single tumor. For clarity, ΔΔG values for ExoD drivers in Group 3 tumors are not shown (they are same as in Group 2). Color in each filled circle-green and shades of green, structure-destabilizing variants (positive ΔΔG); white, variants that are within the standard deviations of ±1.45 kcal/mol and are structure neutral; red and shades of red, structure-stabilizing variants (negative ΔΔG). Yellow, nonsense, or frameshift variants. See **Supplementary Table 6** for more details.

To further analyze *POLE* missense variants in Groups 2, 3, and 4, the AlphaFold2 POLE structure models (**Figure 4A** and **4B**) were used to annotate and analyze each variant by the domain (n=168 variants). A total of 35 destabilizing (ΔΔG ≥ +1.45 kcal/mol) and 17 stabilizing (ΔΔG ≤ −1.45 kcal/mol) missense variants were located at the N-Terminal Lobe (NTL) of the protein in the DNA-unbound model (**Supplementary Table 6**). By domains in the NTL, the N-terminal subdomain (NTD) had 6 destabilizing and 1 stabilizing variants, the ExoD had 8 destabilizing and 4 stabilizing variants (known ExoD drivers excluded), and the polymerase domain had 11 destabilizing and 7 stabilizing variants in the DNA-unbound models (**Supplementary Table 6**). One additional destabilizing and one other stabilizing mutation in the NTL occurs in the unstructured N-terminal segment (residues 1-28) in the DNA-unbound models. By contrast, the C-Terminal Lobe (CTL) had 10 destabilizing and 22 stabilizing variants in the DNA-unbound models (**Supplementary Table 6**). Overall, we found that structure-destabilizing missense variants were more prevalent in the NTL than the CTL (see **Supplementary Table 6;** red = stabilizing; green=destabilizing).

**Figure 4.**
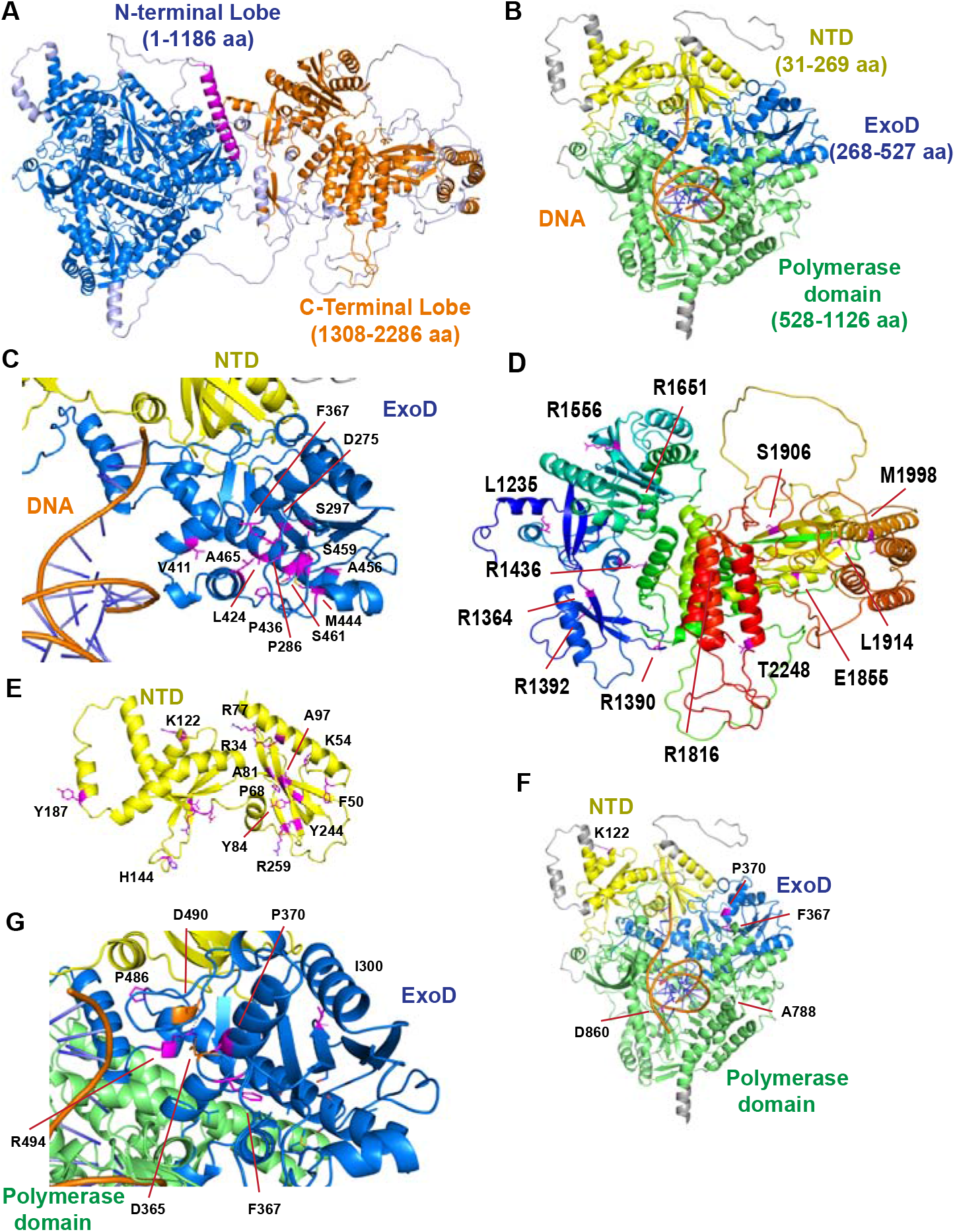
Structure-function assessment of *POLE* variants in Group 2, 3, and 4 tumors (CLS dataset). Human POLE structure models based on yeast POLE templates without DNA bound (full-length yeast POLE with Dbp2, Dbp3, and Dbp4 subunits, PDB:6WJV (21) and with DNA bound (N-terminal lobe only, PDB:4M8O (22) and were generated using AlphaFold2 (18,19). **A**. The length of the POLE protein is 2286 amino acids (aa). The structure can be divided into the NTL (aa 1-1186, blue) and the CTL (aa 1308-2286, orange), which are connected via a linker (aa 1187-1307, mostly weakly predicted, shown in gray, except for an interdomain helix in magenta, aa 1264-1292). **B.** The NTL contains the NTD (aa 31-269, yellow), the ExoD (aa 268-527, blue), and the polymerase domain (aa 528-1186, green). The polymerase domain is further divided into the palm (aa 528-970), the structurally “flexible” part of the palm called the finger (aa 753-833), and the thumb (aa 951-1186) (23,24). The ExoD and polymerase domains together (aa 270-1186) perform DNA synthesis and repair. The roles of the NTD of the NTL and the CTL are not well-studied; the CTL stabilizes the POLE structure and interacts with other subunits in the POLE holoenzyme complex (Dpb2, Dpb3, and Dpb4) (23). **C**. Structural context of the ExoD driver mutations (n=20). None of the residues are in direct contact with DNA, as determined by superposing the AlphaFold2 model based on PDB:4M8O with the yeast POLE/DNA complex structure in PDB:4M8O. **D**. Structural context of the POLE CTL variants. **E.** Structural context of the POLE N-terminal variants (co-occurring with V411L driver). The 7 variants in the V411L + one variants Group 3 tumors associated with higher mTMB are scattered across the full length of POLE (K122N in the NTD, and P370T in the ExoD domain, A788V and D860G in the polymerase domain, R1233* and Q1239R in the linker, and R2131C in the CTD). P370T and D860G in the NTL are destabilizing according to the ddG_monomer. **F, G**. Assessment of variants from the NTD subdomain and the ExoD with striking (highly destabilizing or stabilizing) ΔΔG values (see **Supplementary Text**).

The 20 ExoD driver residues (**Figure 4C, Supplementary Table 6**, drivers defined in **Methods)** are almost all buried in the hydrophobic core of the ExoD, even though they are not all hydrophobic. The most destabilizing driver mutations in both the without-DNA (all >4.0 kcal/mol) and with-DNA (all >2.8 kcal/mol) Rosetta ddG_monomer calculations are driver mutations of hydrophobic amino acids: P286R, L424F, P436S, P436Y, P436R, M444K, A456P. Other structure destabilizing driver mutations of hydrophobic amino acids are F367C, S459Y, and S461P. These drivers are almost all either in the central helix of the ExoD (“Exo III motif”(25)) or in contact with it (**Figure 4C**); implying effect on stability or dynamics of the ExoD. Groups 3 and 4 ExoD variants not known to be drivers that also have highly destabilizing ddG_monomer values include M295R, I300S, F320V, V334G, A480D, and P486S, none of which occur by themselves in tumor samples. A few ExoD drivers were structure stabilizing (S297F/Y, S461L, A465V, S459F) and a few had little impact on predicted stability (F367V/L, V411L).

In Group 3 tumors with P286R + one variant all but one of the 20 mutations at 17 sites have TMB above the median value for P286R alone (mTMB=103) (**Supplementary Table 7**). Thirteen of these 17 sites (77%) are in the CTL which binds accessory proteins (R1364C, R1382C, R1390C, R1436W, R1556W, R1651K, R1826W, E1855D, S1906Y, L1914I, M1998I, T2248I) or the linker between the NTL and CTL (L1235I); five of these are predicted to be in contact with the DNA POLE subunit 2 (L1235I, R1364C, R1390C, R1826W, T2248I), which is present in the template for the model of human POLE without DNA (**Figure 4D**, and **4E**). In Group 3 tumors with V411L driver, there are 7 tumors with V411L + one variant, and they all have TMB above the median value of V411L alone (mTMB=44) (**Figure 4D, 4F** and **Supplementary Table 7**). In contrast to P286R, the seven variants associated with higher mTMB in the presence of V411L are scattered across the length of POLE (**Figure 4D** and **4F)**. **Figure 4G** shows POLE variants from the NTD subdomain and the ExoD domain with striking ΔΔG values, some with the most highly destabilizing ΔΔG values (see **Supplementary Text**).

## Discussion

It is currently not known why tumors of a given cancer type and with the same *POLE* ExoD driver have different levels of TMB, i.e., some show hypermutation (10-100 mut/Mb) while others are ultra-hypermutated (>100 mut/Mb). This study uncovers a distinct subset of highly mutated *POLE* ExoD-mutated tumors with additional *POLE* variants in several folded motifs of POLE. Our findings suggest that a route to acquisition of ultra-hypermutation is through acquisition of one or more additional variants in *POLE* beyond the established ExoD driver. In fact, the 3-nucleotide context of these *POLE* variants suggest that they are secondary to the proofreading defect in those tumors. In the three cancer types studied, CRC, EC, and OC, TMB was significantly higher when the corresponding *POLE* ExoD driver mutation was present in conjunction with one or more additional *POLE* variant(s). This result was still significant when MSI-H tumors were excluded from the analysis (except for OC, likely due to the smaller sample size). These findings were validated in polymerase ε proofreading deficient tumors from TCGA. As observed before (26), tumors with *POLE* ExoD driver mutations were diagnosed earlier than proofreading proficient tumors without a *POLE* ExoD driver. Moreover, we also observed a trend to earlier age of onset for CRCs and OCs with *POLE* ExoD driver occurring in conjunction with other *POLE* variants.

Restricting TMB analysis to specific ExoD hotspot drivers showed that both stronger (e.g., P286R) and weaker (e.g., V411L) ExoD drivers had additional *POLE* variant(s). However, tumors with V411L or other drivers had more *POLE* variants versus tumors with P286R, which at most harbored one additional POLE variant. Recent studies have revealed several differences in proofreading-defective *POLE*-mutated tumors. lt has been shown that all ExoD mutations do not have strong mutagenic effects, and sometimes mutations in the polymerase domain can be associated with hypermutation (10,16). Typically, patient tumor genomic profiles and tumor cell lines do not exhibit evidence of loss of heterozygosity (LOH) for the *POLE* ExoD driver mutation (27). Our data suggest that while inactivation of exonuclease activity is sufficient to drive mutagenesis, it may not always be sufficient to drive ultra-hyper mutagenesis, especially in the case of ExoD drivers with weaker mutagenic effect (e.g., V411L). The three nucleotide sequence contexts of the additional POLE variants in Group 3 tumors strongly suggest that the *POLE* ExoD driver mutation first made the DNA synthesis error, and the cells with the additional *POLE* variant subsequently proliferated and expanded during tumor development. While most additional *POLE* variants were secondary, a minor subset did not fall under the signature sequence contexts associated with POLE defects and could be pre-existing; limitations of retrospective data prevent further analysis on when these mutations emerged. Finally, we found 5 tumors that carried two known ExoD drivers as opposed to a single ExoD driver; limitations of retrospective data prevent further analysis on which ExoD driver was acquired first.

Previously, only simple homology models of human POLE from yeast POLE structures have been used in the structure-function analysis of POLE mutations (11,28). The use of Alphafold2 in this study to model human POLE more accurately allowed us to calculate protein stability changes in the mutant versus the WT proteins to provide unique insights into the known ExoD drivers and POLE variants. Protein stability can more accurately predict changes due to mutations especially missense mutations in proteins (29). We found that while most established ExoD drivers are predicted to be structure destabilizing, there are few established drivers that are either predicted to be structure stabilizing or have no significant predicted impact on stability. For the POLE variants that co-occurred with the ExoD driver, the impact on stability differed by regions or domains; more structure-destabilizing variants were more prevalent in the NTL compared to the CTL. It is possible to speculate that variants in certain regions/domains of POLE (occurring in conjunction with the driver) may lead to other mechanisms increasing mutagenesis beyond simple defects in proofreading. In fact, these other mechanisms beyond simple proofreading defect have already begun to be associated with driver mutations; for example, the P286R mutation is thought to produce a hyperactive DNA polymerization state which amplifies the proofreading defect (30).

This study nominates several fundamental questions that need to be addressed in further studies using biochemical activity assays and/or cellular models to study the secondary *POLE* variants described here in an isogenic genetic background with and without an ExoD driver allele. The data in this study may have implications for clinical management of patients with *POLE*-mutated tumors to understand if there is a systematic clinical benefit associated with tumors carrying specific ExoD drivers plus additional variant(s). Recent studies suggest that this may indeed be the case (11,12,26). Garmezy *et al*. recently reported better clinical outcomes in patients with *POLE* pathogenic variants and in patients with *POLE* VUS (26). Here, most VUS that correlated with better outcomes affected other regions of POLE apart from the ExoD. Rousseau *et al*. found that individuals with tumors bearing *POLE* VUS within the ExoD catalytic site, or the DNA binding site showed clinical benefit from nivolumab (11). Another recent study observed that high TMBs (median: 275.38/Mb) in tumors with POLE proofreading and mismatch repair deficiency were significantly associated both with response to immune checkpoint inhibitors and survival (12). This suggests not all tumors with high TMB may exhibit the same clinical benefit, and that analysis by specific TMB thresholds (which would largely include Group 3 tumors identified in this study) may demonstrate better outcomes.

This study has several limitations: the data are retrospective, and the analysis presented here cannot fully capture the impact of *POLE* variants in conjunction with *POLE* ExoD drivers based on limitations in the dataset on response to therapy, exposures, ancestry/ethnicity. The study cannot always exclude the variants analyzed as germline versus somatic. Additionally, larger sample sizes and longer clinical follow-up studies are needed to investigate the long-term outcomes of patients with such tumors. These data support future mechanistic studies on the synergy of additional *POLE* variants with *POLE* ExoD driver mutations that impact mutagenesis, which is essential for understanding not only tumor development but also patient clinical outcomes.

## Materials and Methods

### Characterization of the discovery cohort (CLS)

We conducted a retrospective analysis on the genomic profiles of 1870 CRC, 4481 EC, and 8190 OC patients that underwent genomic profiling by CLS (Phoenix, AZ) as part of their routine comprehensive tumor molecular profiling. This study was conducted in accordance with 45 CFR 46.101(b), we used retrospective, de-identified patient data and this study was considered IRB exempt. Thereby, no patient consent was necessary from the subjects. All data were obtained through a Data Use Agreement between CLS and Dr. Michael Hall at the Fox Chase Cancer Center (IRB 15-8003).

CLS performed next-generation sequencing on genomic DNA from FFPE tumor samples using the NextSeq platform (Illumina, Inc.). Here, 592 whole-gene targets were enriched using a custom-designed SureSelect XT assay (Agilent Technologies); a total of 1.4 MB was assessed. All reported variants were detected with >99% confidence based on allele frequency and amplicon coverage. The average sequencing depth of coverage was >500 and the analytic sensitivity was of 5%. In the sequencing panel, splice junctions are covered with mutations observed at ±30 nucleotides from the boundaries of *BRCA1/2* genes and ±10 nucleotides of the other genes. Splicing variants were annotated only for mutations detected in ±2 nucleotides from the exon boundaries. The copy-number alteration for each exon was determined as the average depth of the tumor sample along with the sequencing depth of each exon and comparing this data to a precalibrated value. TMB was measured by counting all nonsynonymous missense mutations found per tumor that had not been previously described as germline alterations, and the threshold to define TMB-high was ≥10 mutations/MB; TMB measured by following routinely used guidelines published in (31).

CLS provided patient clinical and demographic data that were collected from electronic medical records between 6/2016 to 6/2019. TMB, tumor lineage, primary tumor site, patient diagnosis, specimen location, age, sex, MSS/MSI status, *POLE* variants, and variants in the 592-targeted gene somatic panel were obtained from CLS. The pathogenicity of each *POLE* mutation was annotated based on the American College of Medical Genetics and Genomics designation as “pathogenic”, “likely pathogenic”, “VUS”, “presumed benign,” or “benign” (32). To stratify patients into groups, we used a list of known *POLE* ExoD drivers (n=20): D275G, P286R, S297F/Y, F367C/L/V, V411L, L424F, P436R/S/Y, M444K/L, A456P, S459F/Y, S461L/P, A465V (10). Total *POLE*-mutated CRC, EC, and OC patient tumor count was n=447 (CRC, n=92; EC, n=307; OC, n=48). TMB threshold of <10 was used to get MSS TMB-L, MSI TMB-L, or TMB≥10 to get MSS/TMB-H, and MSI/TMB-H. The age distribution of the CRC, EC, and OC patients within each of the four groups was used to determine the percentage of frequency of the mutations within each cohort for CRC, EC, OC, and for all the cancers combined. We plotted the age distribution in Graphpad Prism V.9 (https://www.graphpad.com/) and smoothened the curve using Fit Spline (5 knots). See **Supplemental Methods** for additional details.

## Data availability

The deidentified genomic sequencing data are owned by CLS and are not publicly available. The datasets analyzed during the current study are available from the authors upon reasonable request and with permission of CLS. Qualified researchers may contact the corresponding author with request.

## Author contributions

S.S. and E.V.D. performed most studies, data analysis and interpretation, figure, and table generation, writing, and editing; S.R., mutation signatures, TMB analysis by drivers, writing and editing; P.M., J.V-E, & L.V. TCGA analysis and critically reviewed the manuscript; B.F., M.A., and R.L.D. AlphaFold2 models, ddG Rosetta, writing, discussion; J.X., MK. patient data; M.J.H. discussion, designing study, obtained data; S.A., discussion, designing, planning studies, writing & editing.

## Acknowledgments

The Molecular Modeling core facility at the Fox Chase Cancer Center (FCCC) contributed to this study. We express gratitude to Drs. Erica Golemis, Alfonso Bellacosa, Karthik Devarajan, and Adria Hasan at FCCC for their critical input on the article. We also express gratitude to Christine O’Donnell at FCCC for her assistance in preparing the manuscript for publication. The results shown here are in part based upon data generated by TCGA Research Network: https://www.cancer.gov/tcga.

## Abbreviations

CLS: Caris Life Sciences
CRC: Colorectal cancer
CTL: C-terminal Lobe
FCCC: Fox Chase Cancer Center
ΔΔG: change in Gibbs free energy
EC: endometrial cancer
ExoD: exonuclease domain
ICI: immune checkpoint inhibitor
mTMB: median tumor mutational burden
MSI: microsatellite instability
MSS: microsatellite stable
NTL: N-terminal Lobe
NTD: N-terminal subdomain
SBS: single base substitution
POLE: DNA polymerase epsilon
TMB: tumor mutation burden
TMB-H: tumor mutation burden high
TMB-L: tumor mutation burden low
VUS: variants of uncertain significance

## References

1. Nicolas E, Golemis EA, Arora S. POLD1: Central mediator of DNA replication and repair, and implication in cancer and other pathologies. Gene 2016;590(1):128–41 doi 10.1016/j.gene.2016.06.031.

2. Vande Perre P, Siegfried A, Corsini C, Bonnet D, Toulas C, Hamzaoui N, et al. Germline mutation p.N363K in POLE is associated with an increased risk of colorectal cancer and giant cell glioblastoma. Fam Cancer 2019;18(2):173–8 doi 10.1007/s10689-018-0102-6.

3. Mur P, García-Mulero S, Del Valle J, Magraner-Pardo L, Vidal A, Pineda M, et al. Role of POLE and POLD1 in familial cancer. Genet Med 2020;22(12):2089–100 doi 10.1038/s41436-020-0922-2.

4. Palles C, Cazier JB, Howarth KM, Domingo E, Jones AM, Broderick P, et al. Germline mutations affecting the proofreading domains of POLE and POLD1 predispose to colorectal adenomas and carcinomas. Nat Genet 2013;45(2):136–44 doi 10.1038/ng.2503.

5. Valle L, Hernández-Illán E, Bellido F, Aiza G, Castillejo A, Castillejo M-I, et al. New insights into POLE and POLD1 germline mutations in familial colorectal cancer and polyposis. Human Molecular Genetics 2014;23(13):3506–12 doi 10.1093/hmg/ddu058.

6. Djursby M, Madsen MB, Frederiksen JH, Berchtold LA, Therkildsen C, Willemoe GL, et al. New Pathogenic Germline Variants in Very Early Onset and Familial Colorectal Cancer Patients. Frontiers in Genetics 2020;11 doi 10.3389/fgene.2020.566266.

7. Germline DNA Polymerase Mutations Increase Cancer Susceptibility. Cancer Discovery 2013;3(2):136– doi 10.1158/2159-8290.Cd-rw2013-006.

8. Ahn S-M, Ahmad Ansari A, Kim J, Kim D, Chun S-M, Kim J, et al. The somatic POLE P286R mutation defines a unique subclass of colorectal cancer featuring hypermutation, representing a potential genomic biomarker for immunotherapy. Oncotarget 2016;7(42).

9. León-Castillo A, Britton H, McConechy MK, McAlpine JN, Nout R, Kommoss S, et al. Interpretation of somatic POLE mutations in endometrial carcinoma. J Pathol 2020;250(3):323–35 doi 10.1002/path.5372.

10. Campbell BB, Light N, Fabrizio D, Zatzman M, Fuligni F, de Borja R, et al. Comprehensive Analysis of Hypermutation in Human Cancer. Cell 2017;171(5):1042–56.e10 doi 10.1016/j.cell.2017.09.048.

11. Rousseau B, Bieche I, Pasmant E, Hamzaoui N, Leulliot N, Michon L, et al. PD-1 Blockade in Solid Tumors with Defects in Polymerase Epsilon. Cancer Discov 2022;12(6):1435–48 doi 10.1158/2159-8290.Cd-21-0521.

12. Das A, Sudhaman S, Morgenstern D, Coblentz A, Chung J, Stone SC, et al. Genomic predictors of response to PD-1 inhibition in children with germline DNA replication repair deficiency. Nat Med 2022;28(1):125–35 doi 10.1038/s41591-021-01581-6.

13. Chung J, Maruvka YE, Sudhaman S, Kelly J, Haradhvala NJ, Bianchi V, et al. DNA Polymerase and Mismatch Repair Exert Distinct Microsatellite Instability Signatures in Normal and Malignant Human Cells. Cancer Discov 2021;11(5):1176–91 doi 10.1158/2159-8290.Cd-20-0790.

14. Rahn S, Krüger S, Mennrich R, Goebel L, Wesch D, Oberg HH, et al. POLE Score: a comprehensive profiling of programmed death 1 ligand 1 expression in pancreatic ductal adenocarcinoma. Oncotarget 2019;10(16):1572–88 doi 10.18632/oncotarget.26705.

15. Haruma T, Nagasaka T, Nakamura K, Haraga J, Nyuya A, Nishida T, et al. Clinical impact of endometrial cancer stratified by genetic mutational profiles, POLE mutation, and microsatellite instability. PLoS One 2018;13(4):e0195655 doi 10.1371/journal.pone.0195655.

16. Hodel KP, Sun MJS, Ungerleider N, Park VS, Williams LG, Bauer DL, et al. POLE Mutation Spectra Are Shaped by the Mutant Allele Identity, Its Abundance, and Mismatch Repair Status. Mol Cell 2020;78(6):1166–77.e6 doi 10.1016/j.molcel.2020.05.012.

17. Robinson PS, Coorens THH, Palles C, Mitchell E, Abascal F, Olafsson S, et al. Increased somatic mutation burdens in normal human cells due to defective DNA polymerases. Nat Genet 2021;53(10):1434–42 doi 10.1038/s41588-021-00930-y.

18. Jumper J, Evans R, Pritzel A, Green T, Figurnov M, Ronneberger O, et al. Highly accurate protein structure prediction with AlphaFold. Nature 2021;596(7873):583–9 doi 10.1038/s41586-021-03819-2.

19. Tunyasuvunakool K, Adler J, Wu Z, Green T, Zielinski M, Žídek A, et al. Highly accurate protein structure prediction for the human proteome. Nature 2021;596(7873):590–6 doi 10.1038/s41586-021-03828-1.

20. Alford RF, Leaver-Fay A, Jeliazkov JR, O’Meara MJ, DiMaio FP, Park H, et al. The Rosetta All-Atom Energy Function for Macromolecular Modeling and Design. J Chem Theory Comput 2017;13(6):3031–48 doi 10.1021/acs.jctc.7b00125.

21. Yuan Z, Georgescu R, Schauer GD, O’Donnell ME, Li H. Structure of the polymerase ε holoenzyme and atomic model of the leading strand replisome. Nat Commun 2020;11(1):3156 doi 10.1038/s41467-020-16910-5.

22. Hogg M, Osterman P, Bylund GO, Ganai RA, Lundström EB, Sauer-Eriksson AE, et al. Structural basis for processive DNA synthesis by yeast DNA polymerase □. Nat Struct Mol Biol 2014;21(1):49–55 doi 10.1038/nsmb.2712.

23. Yuan Z, Georgescu R, Schauer GD, O’Donnell ME, Li H. Structure of the polymerase ε holoenzyme and atomic model of the leading strand replisome. Nature Communications 2020;11(1):3156 doi 10.1038/s41467-020-16910-5.

24. Baranovskiy AG, Gu J, Babayeva ND, Kurinov I, Pavlov YI, Tahirov TH. Crystal structure of the human Pol□ B-subunit in complex with the C-terminal domain of the catalytic subunit. J Biol Chem 2017;292(38):15717–30 doi 10.1074/jbc.M117.792705.

25. Dahl JM, Thomas N, Tracy MA, Hearn BL, Perera L, Kennedy Scott R, et al. Probing the mechanisms of two exonuclease domain mutators of DNA polymerase □. Nucleic Acids Research 2022;50(2):962–74 doi 10.1093/nar/gkab1255.

26. Garmezy B, Gheeya J, Lin HY, Huang Y, Kim T, Jiang X, et al. Clinical and Molecular Characterization of POLE Mutations as Predictive Biomarkers of Response to Immune Checkpoint Inhibitors in Advanced Cancers. JCO Precis Oncol 2022;6:e2100267 doi 10.1200/po.21.00267.

27. Park VS, Pursell ZF. POLE proofreading defects: Contributions to mutagenesis and cancer. DNA Repair (Amst) 2019;76:50–9 doi 10.1016/j.dnarep.2019.02.007.

28. Parkash V, Kulkarni Y, Ter Beek J, Shcherbakova PV, Kamerlin SCL, Johansson E. Structural consequence of the most frequently recurring cancer-associated substitution in DNA polymerase ε. Nat Commun 2019;10(1):373 doi 10.1038/s41467-018-08114-9.

29. Cao H, Wang J, He L, Qi Y, Zhang JZ. DeepDDG: Predicting the Stability Change of Protein Point Mutations Using Neural Networks. J Chem Inf Model 2019;59(4):1508–14 doi 10.1021/acs.jcim.8b00697.

30. Xing X, Kane DP, Bulock CR, Moore EA, Sharma S, Chabes A, et al. A recurrent cancer-associated substitution in DNA polymerase ε produces a hyperactive enzyme. Nat Commun 2019;10(1):374 doi 10.1038/s41467-018-08145-2.

31. Merino DM, McShane LM, Fabrizio D, Funari V, Chen SJ, White JR, et al. Establishing guidelines to harmonize tumor mutational burden (TMB): in silico assessment of variation in TMB quantification across diagnostic platforms: phase I of the Friends of Cancer Research TMB Harmonization Project. J Immunother Cancer 2020;8(1) doi 10.1136/jitc-2019-000147.

32. Fortuno C, Lee K, Olivier M, Pesaran T, Mai PL, de Andrade KC, et al. Specifications of the ACMG/AMP variant interpretation guidelines for germline TP53 variants. Hum Mutat 2021;42(3):223–36 doi 10.1002/humu.24152.

