## Supplemental Text for "Co-occurring mutations in the POLE exonuclease and non-exonuclease domains define a unique subset of highly mutagenic tumors"

**Supplemental Results.**

**Effect on protein stability of the *POLE* missense variants identified in Groups 2, 3 and 4 tumors.**

Using PyMOL (1) we visually assessed variants from the NTD subdomain and the ExoD domain which had striking ΔΔG values (**Figure 3**, **Supplementary Table 6**). Variants in the NTD were commonly observed (R34H, F50V, K54N, P68S, R77C, A81V, Y84D, A97V, K122N, V134I, H144R, Y187H, R222C, E223D, Y244C, R259C, F274V) (**Figure 4E**, **Supplementary Table 2**). As little is known about the functional roles of the NTD, we suspect that the effects of these variants range from minor stability changes to larger destabilizations that could disrupt the folding of the entire N-terminal Lobe and polymerase and exoD catalysis.

The ExoD variant, R494W is in a small α-helix which is part of the ExoD domain that interacts with DNA (**Figure 4G**). In the structure destabilizing variant, R494W (ΔΔG = 0.65 kcal/mol with DNA), the change from arginine (R) to tryptophan (W) is significant. R494 is predicted to make salt bridges with ExoD residues D365 and E396. Substitution with Trp maintains the hydrogen bond with D365 and packs neatly against hydrophobic residues of the ExoD domain. W494 was found in three EC patients: in one patient with several other variants (R1471C, R1579C, R1626C, A566T, F931fs) and the ExoD driver, F367L (slightly destabilizing with DNA), and in two patients with the ExoD driver P286R (structure destabilizing, +5.1 (with DNA) and +5.6 kcal/mol (without DNA)) (**Figure 4G**, **Supplementary Table 2)**. The neighbor variant, P486S is highly structure-destabilizing (+8.0 and +6.7, with and without DNA) and is in a loop not far from the substrate DNA (**Figure 4G**). The wildtype P486 makes significant contacts with hydrophobic residues that are removed on mutation to serine. The ExoD-DNA binding site variant mentioned above, P370T (predicted to slightly destabilize in a model with DNA, but significantly destabilize in a model without DNA, +4.3 kcal/mol), co-occurred with V411L in an OC patient (**Supplementary Table 2)**. Typically a change from or to a proline is profound because of the restrictions proline places on backbone conformations compared to other amino acids (2). P370 is close to established driver positions: P286R and F367C/L/V, with F367 and P370 in the same α-helix of the ExoD (**Figure 4G**).

Variant M299V (NTD and ExoD, not predicted to have significant stability changes), I300S (ExoD, structure destabilizing) and F274V (NTD, structure destabilizing) are close to E277 which participates in catalysis via coordination of metal ions (3) (**Supplementary Table 6**). M299V is in a β-sheet that is formed by both NTD and ExoD residues (part of a catalytic site, **Figure 4G**). This β-sheet is part of the hydrophobic core, and sequence changes in this region may not only disrupt the hydrophobic core but may directly impact POLE catalytic activity (3). In I300S, the change from isoleucine to serine is adjacent to S300 and T354. This may disrupt the β-sheet folding as indicated by the positive ΔΔG values (**Supplementary Table 6**). F274 has an important structural role in ExoD-NTD β-sheet formation, and the change from phenylalanine to valine (F274V) likely disrupts NTD-exoD β-sheet interaction with the other ExoD regions which may destabilize the entire ExoD domain. A highly destabilizing variant, V334G (**Supplementary Table 6**), was also found in the same β-sheet (**Figure 4G**). Future biochemical and cellular studies will support the impact of these variants.

**_POLE_ variants in MSS TMB-H subset of Group 4 may be potential new drivers.**

In Group 4 tumors, the *POLE* variants were: 27 ExoD variants (2 deletion-insertion, 1 splice site variant, and 24 missense variants) and 127 non-ExoD variants (8 nonsense mutations, 10 frameshift mutations, 5 deletions, 1 duplication, 1 insertion, 2 canonical splice site variants, and 100 missense variants). When segregating by cancer type, *POLE* variants in Group 4 were observed in CRC (n=21), EC (n=118), and in OC (n=4). Among the MSS TMB-H subset of Group 4 tumors, *POLE* variants associated with TMB-H were observed in CRC and EC (n=14); most variants were missense (8/14), but nonsense and other alterations were also observed (6/14). We speculate that the *POLE* variants in MSS tumors with TMB-H could be potential new drivers. Here, six variants were in the CTD (E1376D, R1386W, Q1475X, P1547S, Y1889X, S1930X) and two variants were in the polymerase domain (R680C, palm; R1125X, thumb). ExoD variants (3/14) were observed as missense (M295R (in CRC and EC), F320V, M444I (also reported in (4)), or as chromosomal alterations (F285_P286delinsLR and N423_L424delinsKI). Interestingly, ExoD variant M295R (predicted to significantly destabilize in a model with DNA or without DNA, ΔΔG without DNA +6.653, ΔΔG with DNA +3.89) was observed in two patients (EC and CRC). POLE M295R is in a β-sheet which is part of the catalytic site, with neighboring residue S297 in the ExoD catalytic center(3). Other established ExoD drivers with high destabilizing ΔΔG scores are also in this region (**Supplementary Figure 1B)**. In a CRC patient, M295R co-occurred with CTD S1930X variant, while in an EC patient it co-occurred with CTD R1125X and R1386W variant (**Supplementary Table 2**).

### Supplemental Methods.

#### TCGA validation cohort.

Sequencing data from 46 POLE proofreading deficient TCGA tumor samples (independently of tumor type) and their MSI status (EC, n=30; CRC, n=6; Breast Cancer (BC), n=1; other, n=9), were obtained from cbioportal (5,6). The webtool Single Mutational Signatures in Cancer (MuSiCa) was used to calculate the TMB in those tumors(7). Similar to the discovery cohort, *POLE* alterations and the TMB data were used to segregate the data into either '*POLE* ExoD driver' (n=17) (EC, n=12; CRC, n=3; other, n=2) or '*POLE* ExoD driver plus *POLE* Variant' (n=29) (EC, n=18; CRC, n=3; BC, n=1; other, n=7); these cohorts were compared using a Mann-Whitney statistical test. The analysis was performed by including and excluding data for MSI-H tumors (n=6).

#### Structural Mapping of *POLE* variants of interest.

To understand the possible structural impact of the *POLE* variants, we assessed these variants and POLE drivers through structural modeling onto 3D models of human POLE. Originally, we generated models using traditional template-driven homology modeling (8) (based on yeast POLE structures PDB codes 4M8O (9) and 6WJV (10)), followed by Rosetta FastRelax methods (11-13), side-chain rebuilding with SCWRL4 (14,15), and analysis with UCSF Chimera (16) software. To build a more accurate full-length model of the human protein without missing loops or segments, we utilized the AlphaFold2 program for protein structure prediction.

AlphaFold2 (AF2) has provided reliable protein structure prediction without the previous requirement for templates with close homology (17), and DeepMind has made publicly available models for the whole human proteome as well as nearly all sequences in UniProt (18). The standard AF2 implementation using graphical processor units (GPUs) has a size limit of about 1400 amino acids (19) on 16 Gbyte GPUs available to us. One approach to overcoming this limitation is to produce models of overlapping fragments of 1400 amino acids or less, and then combining them by superposition. This can lead to large steric clashes and impossible models. Instead, we implemented AF2 using parallelized central processing units (CPUs), capable of producing protein models of up to 3000 amino acids on a specialized workstation.

AF2 can be used with or without the use of experimental structures of templates homologous to the query. This is useful when a specific conformational state is required, that might occur, for instance, in the presence or absence of DNA or accessory proteins. Thus, our implementation of AF2 allows us to choose one or more specific templates rather than searching the entire PDB for all relevant templates. There are several structures of yeast POLE with and without DNA and with various accessory proteins. These structures can be used as templates in AF2. We (and others) have found through trial and error that shallower multiple sequence alignments (MSAs) allow AF2 to adhere more closely to the templates it is given. Our implementation allows us to provide a set of protein sequences as the search database for creating the MSA and a list of one or more templates for modeling proteins in specific states observed in the chosen templates.

With the advent of many new cryo-EM structures of functional multi-protein complexes (i.e., polymerase holoenzymes), there is a need to predict the structure of a protein of interest in a conformation that accounts for the presence of other accessory proteins required for holoenzyme function. In this specific case we used our AlphaFold2 implementation to predict the structure of human POLE and bias the model conformation to account for the other accessory subunits observed in the yeast holoenzyme cryo-EM structure, comprising yeast DNA polymerase epsilon subunits A, B, C, and D (PDB code 6WJV (10)). AlphaFold2 at the EBI database predicts the human POLE model with the C-terminal half of the protein (termed C-Lobe) in a completely different orientation. While the structure and function of the C-Lobe is understudied compared to the details known about the polymerase and exoD portions contained in the N-terminal half, the scaffolding role of the C-Lobe in binding the accessory factors is revealed in the yeast holoenzyme structure. Our AF2 implementation allows an adjustment of the relative weights of a homologous template structure and the multiple sequence alignment that contribute to the final models predicted. With such adjustments, we were able to produce a model of human POLE that aligns well with the yeast homolog with an RMSD of 1.02 $Å^2$ for 1002 alpha-carbon pairs in the NTD half, as well as the linker and C-Lobe found in a similar position seen in the template structure (PDB:6WJV).

As the yeast protein is known to undergo a significant conformational change upon binding to DNA in two major alpha helices in the finger region, we also generated an AF2 model of the N-terminal half of human POLE based on the structure of the yeast POLE crystallized in the presence of a primer-template DNA molecule (PDB code 4M8O (9)). This model was subjected to the same ΔΔG_monomer process to score all the variants that are located in the N-Lobe portion of POLE (amino acids 1 to 1183).

To assess quantitatively the effects of missense mutations in POLE, we employed applications found in the Rosetta Molecular Modeling Suite (11,20). The application named ddG_monomer makes the substitution encoded by the variants and allows repacking of side chains with backbone movement permitted when needed for optimal packing without clashes. As a preparation for using this application, we subjected the AF2 POLE models to Rosetta energy minimization with constraints on coordinate movement (min_cst setting of 0.5Å). The ddG_monomer application generated 25 output structures ("decoys") of the wildtype and each variant and the Rosetta scoring application (2016 energy terms with the jd2score application) generates the ΔG of folding for each decoy. The average ΔG of the 25 decoys for each variant is compared the average ΔG for the wildtype residue yielding a ΔΔG for each variant (abbreviated ΔΔG).

The resulting ΔΔG is given in Rosetta energy units, which have been calibrated to be equivalent to kcal/mol units (21). Mutants with negative ΔΔG's are predicted to be more stable than the wildtype; mutants with positive ΔΔG's are destabilized relative to the wildtype residue. It should be noted that this calculated change in stability does not always correlate with effects on protein function. These cautions notwithstanding, we set out to calculate the ΔΔG's for more than 170 variants in human POLE found in CRC, EC, and OC (see **Supplemental Table 6**). For mutations in the N-terminal lobe, we ran calculations with ddG_monomer with both models and refer to these as "with DNA" (based on PDB:4M8O) and "without DNA" (based on PDB:6WJV). In both cases, the ddG_monomer calculations are performed on the model POLE only, and do not include the DNA or B, C, and D subunits of polymerase epsilon. We used these values to help us judge how each variant would be tolerated in the nearby chemical environment and predict the effect each substitution might have on POLE function. Consideration was given to the sub-domain fold and function, and important bonding interactions that might be perturbed, consequently highlighting the likelihood of detriment to various protein functions.

We ran calculations for the mutations in batches and for each run, a separate WT calculation was performed. The WT values are highly reproducible: for instance, the standard deviation of the mean energies for 5 runs without DNA was only 0.16 kcal/mol. To derive a cutoff value for significant stabilization or destabilization, we calculated the standard deviation for each set of 25 decoys for each mutation and the wildtype both with and without DNA. The mean standard deviation across all decoy sets was sigma=1.62 kcal/mol (range 0.92,3.22). For calculations with DNA, the sigma values were 1.83 and 1.79 for WT and mutant decoy sets respectively. The standard deviation of the difference of means (mutant – wildtype) is SD=sqrt(sigma_mut**2/nmut + sigma_wt**2/nmut), where nmut=25. SD for this case is 0.51 kcal/mol. We used a cutoff value of 1.45 kcal/mol, which is recommended for the ddG_monomer application. This is 2.8 standard deviations, so we have labeled only highly destabilizing or stabilizing mutations as significant. The sigma values for the calculations with DNA were smaller (~1.3 kcal/mol) so the cutoff of 1.45 is still reasonable (SD=0.37; 3.9SD=1.45).

***POLE* variant three nucleotide sequence context analysis.**

For tumors bearing a single *POLE* driver and *POLE* Variant, the mutations in the *POLE* variants were assessed using mutational signatures. All *POLE* ExoD driver associated COSMIC mutational signatures (SBS 10a, SBS 10b, SBS 14, and SBS 28) were assessed (22). Each of these signatures has a primary mutation which has been described as a "hotspot" (22); SBS 10a is C>A in TCT context; SBS 10b is C>T in the TCG context; SBS 14 is C>A in the NCT context (N is any base); and SBS 28 is T>G in the TTT context. In addition to these primary “hotspots”, all mutations that constitute >1% of the genome signature of interest were counted, capturing 88-89% of each signature in the analysis.

**Statistical Analysis.**

We used descriptive statistics to determine medians and quartiles for age distributions. The median TMB (mTMB) with range was reported for each individual cohort for each cancer type (CRC, EC, and OC) for the Caris Life Sciences dataset. We combined data from multiple cancer types, where appropriate, to report mTMB with range in MSI-H and MSS or MSI-Low tumors. We used Mann-Whitney tests, where appropriate, to determine the clinical significance of the mTMB difference between the compared cohorts; ***, $p<0.001$; **, $p<0.01$; *, $p<0.05$; NS, non-significant. Corrections for multiple testing were performed for more than two comparisons using the Benjamini Hochberg False Discovery Rate (FDR) test.

**References.**


1. PYMOL. The PyMOL Molecular Graphics System, Version 2.0 (Schrödinger, LLC).
2. Ganguly HK, Basu G. Conformational landscape of substituted prolines. Biophys Rev **2020**;12(1):25-39 doi 10.1007/s12551-020-00621-8.
3. Rousseau B, Bieche I, Pasmant E, Hamzaoui N, Leulliot N, Michon L*, et al.* PD-1 Blockade in Solid Tumors with Defects in Polymerase Epsilon. Cancer Discov **2022**;12(6):1435-48 doi 10.1158/2159-8290.Cd-21-0521.
4. Parra-Herran C, Lerner-Ellis J, Xu B, Khalouei S, Bassiouny D, Cesari M*, et al.* Molecular-based classification algorithm for endometrial carcinoma categorizes ovarian endometrioid carcinoma into prognostically significant groups. Mod Pathol **2017**;30(12):1748-59 doi 10.1038/modpathol.2017.81.
5. Cerami E, Gao J, Dogrusoz U, Gross BE, Sumer SO, Aksoy BA*, et al.* The cBio cancer genomics portal: an open platform for exploring multidimensional cancer genomics data. Cancer Discov **2012**;2(5):401-4 doi 10.1158/2159-8290.Cd-12-0095.
6. Gao J, Aksoy BA, Dogrusoz U, Dresdner G, Gross B, Sumer SO*, et al.* Integrative analysis of complex cancer genomics and clinical profiles using the cBioPortal. Sci Signal **2013**;6(269):pl1 doi 10.1126/scisignal.2004088.
7. Díaz-Gay M, Vila-Casadesús M, Franch-Expósito S, Hernández-Illán E, Lozano JJ, Castellví-Bel S. Mutational Signatures in Cancer (MuSiCa): a web application to implement mutational signatures analysis in cancer samples. BMC Bioinformatics **2018**;19(1):224 doi 10.1186/s12859-018-2234-y.
8. Demidova EV SI, Vlasenkova R, Kelow S, Andrake MD, Hartman TR, Kent T, Pomerantz RT, Dunbrack Jr. RL, Golemis EA, Hall MJ, Chen DYT, Daly MB, Arora S. Signatures of defective DNA repair and replication in early-onset renal cancer patients referred for germline genetic testing. Medrxiv **2022** doi https://doi.org/10.1101/2022.05.23.22275227.
9. Hogg M, Osterman P, Bylund GO, Ganai RA, Lundström EB, Sauer-Eriksson AE*, et al.* Structural basis for processive DNA synthesis by yeast DNA polymerase ε. Nat Struct Mol Biol **2014**;21(1):49-55 doi 10.1038/nsmb.2712.
10. Yuan Z, Georgescu R, Schauer GD, O’Donnell ME, Li H. Structure of the polymerase ε holoenzyme and atomic model of the leading strand replisome. Nature Communications **2020**;11(1):3156 doi 10.1038/s41467-020-16910-5.
11. Alford RF, Leaver-Fay A, Jeliazkov JR, O'Meara MJ, DiMaio FP, Park H*, et al.* The Rosetta All-Atom Energy Function for Macromolecular Modeling and Design. J Chem Theory Comput **2017**;13(6):3031-48 doi 10.1021/acs.jctc.7b00125.
12. Khatib F, Cooper S, Tyka MD, Xu K, Makedon I, Popovic Z*, et al.* Algorithm discovery by protein folding game players. Proc Natl Acad Sci U S A **2011**;108(47):18949-53 doi 10.1073/pnas.1115898108.
13. Maguire JB, Haddox HK, Strickland D, Halabiya SF, Coventry B, Griffin JR*, et al.* Perturbing the energy landscape for improved packing during computational protein design. Proteins **2021**;89(4):436-49 doi 10.1002/prot.26030.

14. Krivov GG, Shapovalov MV, Dunbrack RL, Jr. Improved prediction of protein side-chain conformations with SCWRL4. Proteins **2009**;77(4):778-95 doi 10.1002/prot.22488.
15. Shapovalov MV, Dunbrack RL, Jr. A smoothed backbone-dependent rotamer library for proteins derived from adaptive kernel density estimates and regressions. Structure **2011**;19(6):844-58 doi 10.1016/j.str.2011.03.019.
16. Pettersen EF, Goddard TD, Huang CC, Couch GS, Greenblatt DM, Meng EC*, et al.* UCSF Chimera--a visualization system for exploratory research and analysis. J Comput Chem **2004**;25(13):1605-12 doi 10.1002/jcc.20084.
17. Jumper J, Evans R, Pritzel A, Green T, Figurnov M, Ronneberger O*, et al.* Highly accurate protein structure prediction with AlphaFold. Nature **2021**;596(7873):583-9 doi 10.1038/s41586-021-03819-2.
18. Tunyasuvunakool K, Adler J, Wu Z, Green T, Zielinski M, Žídek A*, et al.* Highly accurate protein structure prediction for the human proteome. Nature **2021**;596(7873):590-6 doi 10.1038/s41586-021-03828-1.
19. Mirdita M, Schütze K, Moriwaki Y, Heo L, Ovchinnikov S, Steinegger M. ColabFold: making protein folding accessible to all. Nat Methods **2022**;19(6):679-82 doi 10.1038/s41592-022-01488-1.
20. Leaver-Fay A, O'Meara MJ, Tyka M, Jacak R, Song Y, Kellogg EH*, et al.* Scientific benchmarks for guiding macromolecular energy function improvement. Methods Enzymol **2013**;523:109-43 doi 10.1016/b978-0-12-394292-0.00006-0.
21. Alford RF, Leaver-Fay A, Jeliazkov JR, O'Meara MJ, DiMaio FP, Park H*, et al.* The Rosetta All-Atom Energy Function for Macromolecular Modeling and Design. J Chem Theory Comput **2017**;13(6):3031-48 doi 10.1021/acs.jctc.7b00125.
22. Hodel KP, Sun MJS, Ungerleider N, Park VS, Williams LG, Bauer DL*, et al.* POLE Mutation Spectra Are Shaped by the Mutant Allele Identity, Its Abundance, and Mismatch Repair Status. Mol Cell **2020**;78(6):1166-77.e6 doi 10.1016/j.molcel.2020.05.012.

**Supplementary Figure Legends**

**Supplementary Figure 1. Overall three nucleotide sequence context of *POLE* variants in Group 3 tumors**. All COSMIC mutational signatures associated with POLE ExoD driver defects (SBS 10a, SBS 10b, SBS 14, and SBS 28) were assessed (22). Pie chart distribution of SBS 10a, SBS 10b, SBS 14, and SBS 28 in CRC**,** EC**,** and OC are combined. Each of these signatures has a primary mutation which has been described as a "hotspot" (22); SBS 10a is C>A in TCT context; SBS 10b is C>T in the TCG context; SBS 14 is C>A in the NCT context (N is any base); and SBS 28 is T>G in the TTT context. In addition to these primary "hotspots", all mutations that comprise >1% of the genome signature of interest were counted, capturing 88-90% of each signature in the analysis. **B. *POLE* variants in Group 4 MSS TMB-H subset**. *POLE* variants found in MSS CRC (round symbol) and EC (triangle symbol) tumor profiles. *POLE* variants are missense (in green), nonsense (in lavender), and any other (in fuchsia), TMB data, in bold in brackets for each *POLE* variant. The annotated POLE regions are NTD (grey, 31-281), ExoD (wheat, 282-527), polymerase (pink, palm: 528-950; cyan, fingers: 769-833; green, thumb: 951-1186), and CTL (dark grey, 1308-2222).

**Supplementary Figure 2. A and B. Comparison of mTMB and ΔΔG values in Group 2 and 3 tumors in Caris dataset.** With AlphaFold2 DNA bound model and Rosetta ddG_monomer, we generated 25 repacked decoys for each mutation and compared the average energy score for these decoys to an average for 25 decoys of the wildtype protein. We used a cutoff of ±1.45 kcal/mol for significant ΔΔG, corresponding to ~2 standard deviations of the differences of the mean Rosetta scores for WT and mutant structures. **A. Comparison of mTMB and ΔΔG values in Group 2 and 3 tumors by the number of *POLE* Variants**. Data for CRC, EC, and OC genomic profiles were combined and ΔΔG values are plotted against the mTMB. For Group 2 or the Group 3 data with + 1 *POLE* variant plots, each filled round circle represents a single tumor genomic profile. **B. Comparison of mTMB and ΔΔG values in Group 2 and 3 tumors by *POLE* ExoD driver.** data for CRC, EC, and OC genomic profiles were combined and ΔΔG values are plotted against the mTMB. **A and B**. Group 3 tumors with multiple variants, a circle next to another circle (without any space) represents a single tumor. For clarity, ΔΔG values for ExoD drivers in Group 3 tumors are not shown (they are same as in Group 2). Color in each filled circle- green or shades of green, structure-destabilizing variants (positive ΔΔG); white, variants that are within the standard deviations of ±1.45 kcal/mol and are structure neutral; red or shades of red, structure-stabilizing variants (negative ΔΔG). Yellow, nonsense, or frameshift variants; Black, ΔΔG values not calculated in the with DNA model. See **Supplementary Table 6** for more details.

**Supplementary Tables.**

**Supplementary Table 1.** *POLE* variant groups, patient clinical and demographic characteristics for the CARIS cohort.

**Supplementary Table 2.** De-identified patient clinical data with nucleotide contexts of the *POLE* variants (excel file).

**Supplementary Table 3.** Age Distribution of CRC, EC, and OC patients with *POLE*-mutated tumors.

**Supplementary Table 4.** mTMB comparisons in the Caris Life Sciences dataset.

**Supplementary Table 5.** mTMB comparisons in TCGA dataset.

**Supplementary Table 6.** Rosetta ΔΔG values for POLE variants and drivers with or without DNA (excel file).

**Supplementary Table 7.** Mutations in Group 3 tumors with P286R or V411L plus one variant and mTMB comparisons.

**Supplementary Table 1.** *POLE* variant groups, patient clinical and demographic characteristics.

| Clinico-pathological factors | *POLE* variant TMB-L (Group 1) | *POLE* ExoD driver TMB-H | | *POLE* Variant TMB-H (Group 4) |
|---|---|---|---|---|
| | | *POLE* ExoD driver (Group 2) | *POLE* ExoD driver + *POLE* Variant (Group 3) | |
| | **Colorectal Cancer** | | | |
| No. of patients | 36/92 | 11/92 | 24/92 | 21/92 |
| mTMB | 6.0(3-9) | 115(61-216) | 264.5(114-414) | 33(10-461) |
| MSS | 36 | 11 | 21 | 11 |
| MSI | 0 | 0 | 3 | 10 |
| <50 yrs | 6 | 5 | 14 | 8 |
| ≥50 yrs | 30 | 6 | 10 | 13 |
| Female | 19 | 1 | 6 | 8 |
| Male | 17 | 10 | 18 | 13 |
| | **Endometrial Cancer** | | | |
| No. of patients | 95/307 | 37/307 | 57/307 | 118/307 |
| mTMB | 7(3-9) | 52(21-314) | 219(53-520) | 17.50(10-273) |
| MSS | 90 | 35 | 41 | 23 |
| MSI | 5 | 2 | 16 | 95 |
| <50 yrs | 6 | 6 | 10 | 5 |
| ≥50 yrs | 89 | 31 | 47 | 113 |
| | **Ovarian Cancer** | | | |
| No. of patients | 24/48 | 12/48 | 8/48 | 4/48 |
| mTMB | 5.0(4-9) | 69(31-379) | 145(51-394) | 14.50(10-21) |
| MSS | 24 | 12 | 6 | 2 |
| MSI | 0 | 0 | 2 | 2 |
| <50 yrs | 4 | 4 | 5 | 3 |
| ≥50 yrs | 20 | 8 | 3 | 1 |

MMR, mismatch repair; MSI, microsatellite instability; MSS, microsatellite stability; mTMB, median tumor mutational burden.

**Supplementary Table 3.** Age Distribution of CRC, EC, and OC patients with *POLE*-mutated tumors.

| | Age Distribution of Patients | | | | | |
|---|---|---|---|---|---|---|
| | Q0 | Q1 | Median | Q3 | Q4 | Mean |
| | **Colorectal Cancer** | | | | | |
| Group 1 | 28 | 52 | 56 | 63.75 | 84 | 57.72 |
| Group 2 | 24 | 40 | 53 | 62 | 69 | 49.64 |
| Group 3 | 28 | 40 | 45.5 | 57.75 | 73 | 48.08 |
| Group 4 | 25 | 37.5 | 56 | 66.5 | 76 | 52.67 |
| | **Endometrial Cancer** | | | | | |
| Group 1 | 22 | 58 | 62 | 70 | 87 | 63.46 |
| Group 2 | 37 | 52 | 57 | 66 | 88 | 58.81 |
| Group 3 | 34 | 52.5 | 58 | 64 | 84 | 58.44 |
| Group 4 | 31 | 59.75 | 66 | 70.25 | 88 | 65.47 |
| | **Ovarian Cancer** | | | | | |
| Group 1 | 44 | 57.25 | 62.50 | 72.25 | 83 | 63.08 |
| Group 2 | 32 | 43 | 51.5 | 55.25 | 59 | 49.42 |
| Group 3 | 34 | 43.5 | 48.5 | 57.75 | 63 | 49.38 |
| Group 4 | 38 | 40.5 | 48 | 51 | 52 | 46.50 |
| | **All Cancers** | | | | | |
| Group 1 | 22 | 57 | 62 | 70 | 87 | 62.07 |
| Group 2 | 24 | 49.25 | 55.5 | 62 | 88 | 55.25 |
| Group 3 | 28 | 47.5 | 55 | 62 | 84 | 54.83 |
| Group 4 | 25 | 58 | 65 | 70 | 88 | 63.06 |

**Supplementary Table 4.** mTMB comparisons in the Caris Life Sciences dataset.

| **Caris data set** | **Group 1: *POLE* variant TMB-L** | | | **Group 2: *POLE* ExoD driver** | | | **Group 3: *POLE* ExoD driver + *POLE* Variant** | | |
|---|---|---|---|---|---|---|---|---|---|
| Cancer type | **CRC** | **EC** | **OC** | **CRC** | **EC** | **OC** | **CRC** | **EC** | **OC** |
| mTMB (range), including MSI & MSS | 6.0 (3-9) | 7 (3-9) | 5 (4-9) | 115 (61-216) | 52 (21-314) | 69 (31-379) | 264.5 (114-414) | 219 (53-520) | 145 (51-394) |
| Statistics | *** | *** | *** | | | | *** | *** | * |
| mTMB (range), excluding MSI | 6.0 (3-9) | 7 (3-9) | 5 (4-9) | 115 (61-216) | 52 (21-301) | 69 (31-379) | 259 (114-414) | 181 (53-520) | 131.5 (51-261) |
| Statistics | *** | *** | *** | | | | *** | *** | NS |

CRC, colorectal cancer; EC, endometrial cancer; OC, ovarian cancer.
*** represents p<0.001 obtained from Mann-Whitney test. * is p<0.05, *** is p<0.001. NS is not significant.
Group 2 was compared with Group 1 and Group 3.

**Supplementary Table 5.** mTMB comparisons in TCGA dataset.

| **TCGA data set** | **Group 2: *POLE* ExoD driver** | **Group 3: *POLE* ExoD driver + *POLE* Variant** |
|---|---|---|
| mTMB (range), n including MSI & MSS | 104.6 (28.2-302.9), 17 | 250.4 (58.3-532.7), 29 |
| Statistics | | *** |
| mTMB (range), excluding MSI | 108.7(28.2-302.9), 16 | 272.1 (58.3-532.7), 24 |
| Statistics | | *** |

*** is p<0.001 obtained from Mann Whitney test.

**Supplementary Table 7.** Mutations in Group 3 tumors with P286R or V411L plus one variant and mTMB comparisons.

| Mutations in Group 3 tumors with P286R + one Variant | mTMB |
|---|---|
| E1855D | 129 |
| F990C | 261 |
| L1235I | 197 |
| L1914I | 156 |
| M1998I | 85 |
| R1364C | 425 |
| R1382C | 301 |
| R1390C | 243 |
| R1436W | 146 |
| R1556W | 119 |
| R1651K | 125 |
| R1826W | 243 |
| R1826W | 175 |
| R2017C | 127 |
| R494W | 167 |
| R494W | 167 |
| R77C | 302 |
| S1906Y | 226 |
| S1906Y | 193 |
| T2248I | 216 |

| Mutations in Group 3 tumors with V411L + one Variant | mTMB |
|---|---|
| A788V | 134 |
| D860G | 394 |
| K122N | 142 |
| P370T | 72 |
| Q1239R | 61 |
| R1233* | 132 |
| R2131C | 173 |

mTMB, median Tumor Mutation Burden value.

**A**

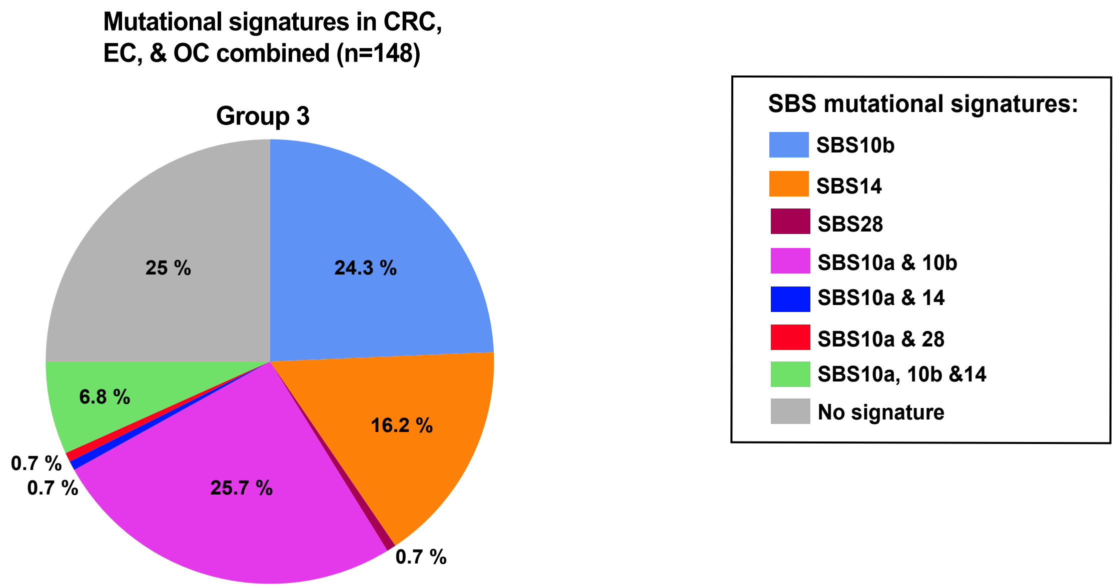


**B**

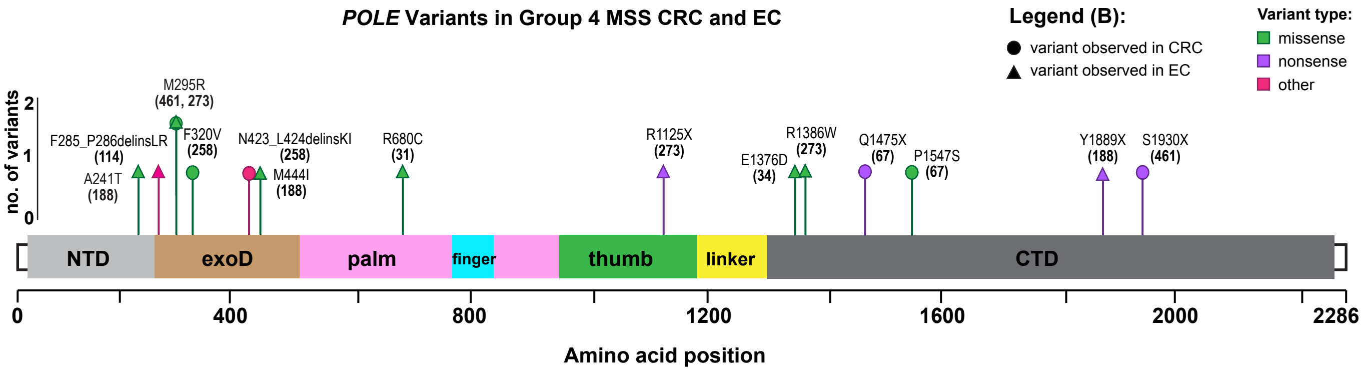


Shah & Demidova et al, Supplementary Figure 1

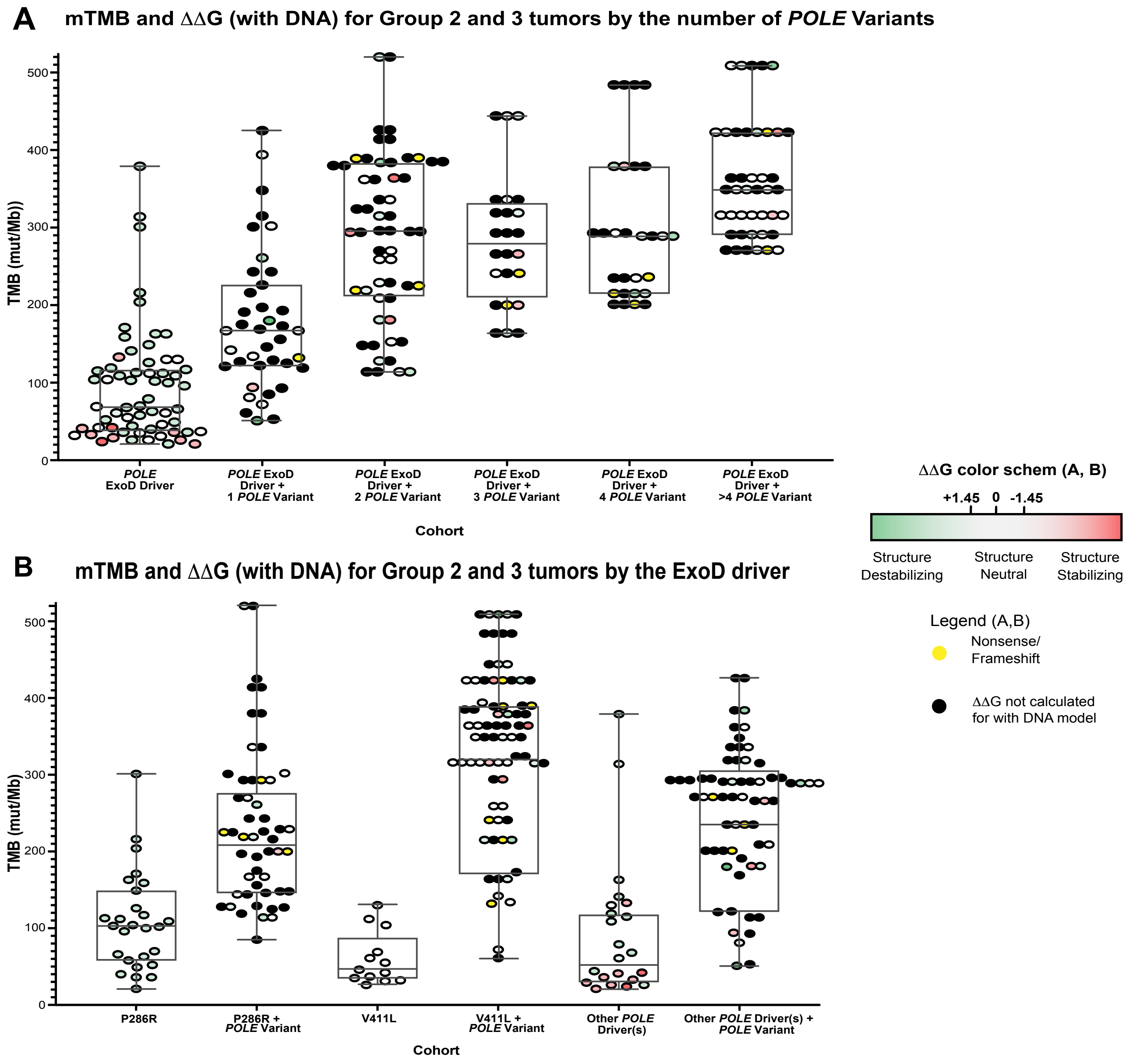


**Shah & Demidova et al., Supplementary Figure 2**
